# Regulatory interactions between APOBEC3B N- and C-terminal domains

**DOI:** 10.1101/2024.12.11.628032

**Authors:** Mac Kevin E. Braza, Özlem Demir, Surl-Hee Ahn, Clare K. Morris, Carla Calvó-Tusell, Kelly L. McGuire, Bárbara de la Peña Avalos, Michael A. Carpenter, Yanjun Chen, Lorenzo Casalino, Hideki Aihara, Mark A. Herzik, Reuben S. Harris, Rommie E. Amaro

**Author notes:** Corresponding Author’s.

## Abstract

APOBEC3B (A3B) is implicated in DNA mutations that facilitate tumor evolution. Although structures of its individual N- and C-terminal domains (NTD and CTD) have been resolved through X-ray crystallography, the full-length A3B (fl-A3B) structure remains elusive, limiting understanding of its dynamics and mechanisms. In particular, the APOBEC3B C-terminal domain (A3Bctd) active site is frequently closed in models and structures. In this study, we built several new models of fl-A3B using integrative structural biology methods and selected a top model for further dynamical investigation. We compared dynamics of the truncated (A3Bctd) to the fl-A3B via conventional and Gaussian accelerated molecular dynamics (MD) simulations. Subsequently, we employed weighted ensemble methods to explore the fl-A3B active site opening mechanism, finding that interactions at the NTD-CTD interface enhance the opening frequency of the fl-A3B active site. Our findings shed light on the structural dynamics of fl-A3B, which may offer new avenues for therapeutic intervention in cancer.

## Introduction

The Apolipoprotein B mRNA editing enzyme, catalytic polypeptide-like 3 (APOBEC3) enzymes are key for human immunity against retroviruses and transposons.^1,2^ APOBEC3 enzymes catalyze the deamination of cytosines to uracil (C-to-U), targeting newly synthesized viral single-stranded DNA (ssDNA). The APOBEC3 family consists of seven cytosine deaminases namely APOBEC3A, APOBEC3B, APOBEC3C, APOBEC3D, APOBEC3F, APOBEC3G, and APOBEC3H. The presence of an APOBEC3-attributed mutational signature (single-base-substitution C-to-T and C-to-G mutations) in the 5’-TC motif in a wide array of cancer types suggests that many of the cancers were associated with A3 activity. This mutagenesis implies their importance in the mutational milieu of human cancer.^3,4^ APOBEC3B (A3B), in particular, is predominantly localized in the nucleus and has been linked to a wide range of cancers, including breast, head, neck, cervical, and bladder cancers, where its expression correlates with drug resistance and metastasis.^5–7^ Furthermore, A3B expression is associated with poor clinical outcomes in estrogen-positive breast cancer patients.^4,8,9^

A3B is one of the double-domain-A3s that has a noncatalytic N-terminal domain (NTD) (sometimes called the “pseudocatalytic” domain) and a catalytic C-terminal domain. Similar to other A3s, A3B chelates one Zn^2+^ ion as a cofactor, bridging the active site residues to the DNA substrate. In recent years, X-ray crystal structures of A3Bctd mutants have been determined.^10–12^ Despite progress in determining the X-ray crystal structures of various A3Bctd mutants, the structure of wild-type fl-A3B remains unresolved, largely due to is tendency to aggregate in solution and poor protein solubility.^13,14^ Although A3B has been proposed as a promising drug target, there is limited knowledge of the active site accessibility and its opening mechanism due to the aforementioned experimental challenges.^10,15,16^

Previous studies have demonstrated that the NTD is involved in DNA and RNA binding and nuclear localization.^7,13,17–19^ Notably, fl-A3B exhibits up to 40-fold greater activity than A3Bctd.^20^ Despite the NTD’s known regulatory role in fl-A3B, there is limited knowledge of its functional significance in the opening of active site and ssDNA deamination activity. Moreover, the absence of fl-A3B structural and dynamical studies limits our understanding of its specificity and mechanism. We hypothesize that the absence of the N-terminal domain (NTD), despite being non-catalytic, might hinder or affect the opening of the active site.

One of the most interesting dynamical properties of A3B is the active site opening. Experimental and computational studies have revealed how rarely the A3Bctd active site opens.^15,19,21^ The A3Bctd active site favors a closed active site conformation and only opened after heavily mutating the active site residues. Furthermore, it remains elusive how the ssDNA binds to A3B active site especially in its wild-type form. We hypothesized that, (experimentally) the absence of NTD and (computationally) the limitation of classical MD simulations hindered the observation of A3B active site opening.

In this work, we used integrative structural biology methods to construct an all-atom model of the fl-A3B and performed extensive all-atom molecular dynamics (MD) simulations to elucidate the role of the NTD in the active site opening. By means of state-of-the-art enhanced sampling methods for MD simulations, such as Gaussian accelerated MD (GaMD) and weighted ensemble (WE) simulations, we characterize the opening mechanism of the ssDNA binding pocket. We find that opening of the fl-A3B active site depends on interactions between residues at the interface of the NTD and the CTD, highlighting the importance of the NTD for more than just DNA binding. This work provides critical insights into the conformational dynamics of fl-A3B, advancing our understanding of the molecular basis underlying ssDNA binding site opening.

## Results and Discussion

In this study, we aimed to elucidate key structural features of full-length APOBEC3B (fl-A3B) that facilitate the opening of the active site. Despite fl-A3B’s significant role in immunity and cancer mutagenesis, experimental structural studies are limited due to inherent solubility and aggregation issues.^13,14^ Furthermore, the specific mechanisms by which the NTD influences nucleic acid recognition and overall protein dynamics remain poorly understood.^10,14,20,23^ Due to extensive evidence that the NTD of other double domain APOBEC3 plays an important role in substrate recognition, we aimed to investigate if the A3B NTD served a similar functional purpose.

### Constructing the full-length APOBEC3B model

ln the absence of an experimentally-resolved fl-A3B structure, homology modeling tools were used to develop full-length models. Different tools including Schrödinger Prime,^24^ I-TASSER,^25^ SWISS-MODEL,^26^ and AlphaFold 2^27^ were used to explore the orientation of the fl-A3B NTD and CTD. The sequence from UniProt corresponding to the human fl-A3B was used and submitted to the software package or web server. As a basis of the NTD and CTD orientation to each other, we used ClusPro web server to generate protein-protein docked poses of two domains.^28^ Surveying the results of the various modeling tools, differences in the predicted orientation of the NTD and CTD were apparent. AlphaFold models exhibited poor model confidence for the loop regions, most especially the active site loops 1, 3, and 7 (Figure 1). AlphaFold 2 and AlphaFold 3 exhibited low predicted local distance difference test (pLDDT) scores at critical active site loops, potentially impacting the accuracy of conformational dynamics. The models developed here were superimposed alongside the experimentally-characterized functionally-related double-domain deaminase, A3G.^22^

**Figure 1.**
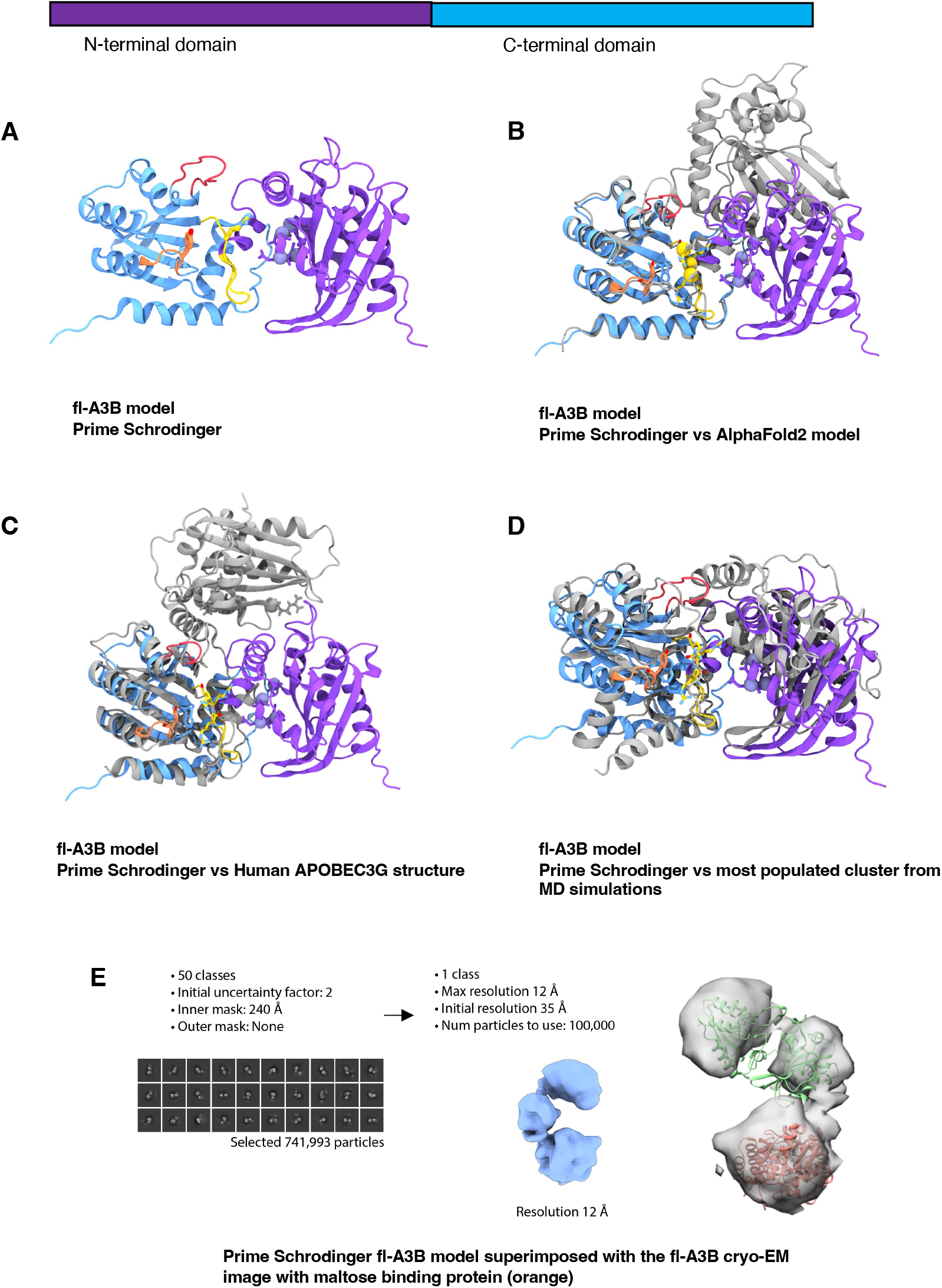
Comparative modeling of fl-A3B. (a) fl-A3B model from Prime - Schrodinger superimposed to (b) AlphaFold2 model; (c) Cryo-EM structure of wild-type human APOBEC3G (PDB ID 8CX0)^22^, (d) most populated clustered structure from our MD simulations; and (e) Representative particle from the cryo-EM experiments of the fl-A3B and maltose binding protein complex.

To attempt to characterize the full-length structure, we performed cryo-EM experiments of fl-A3B bound to maltose binding protein (Figure 1E, SI Figure 1). This enabled the characterization of the fl-A3B structure to maximal resolution of 12 Å. We subsequently aligned all predicted models (Figure 1) to the cryo-EM density. Ultimately, the fl-A3B model from Schrödinger Prime achieved the best (and substantial) alignment with low-resolution cryo-EM particles. We thus selected the Schrödinger Prime model for subsequent MD simulations.

### Full-length APOBEC3B active site opens more frequently than the APOBEC3B C-terminal domain alone

After modeling both the A3Bctd and fl-A3B, we ran a series of conventional MD simulations to characterize their baseline dynamics. Here, we used AMBER pmemd.cuda^29^ with ff14SB^30^ and TIP3P^31^ force fields for protein and water, respectively. Our group and several others have shown that the A3Bctd’s active site (WT and mutated) remain closed in the absence of ssDNA substrate.^15,18,19,21,32,33^ Experimentally, it has been demonstrated that residues R211 and Y315 are crucial for ssDNA substrate interaction within A3B. R211, located in loop 1, interacts with the -1 thymine nucleotide of ssDNA, while Y315 in loop 7 plays a critical role in the active site opening.^10^ Notably, the distances between the N atom of R211 and the OH group of Y315 differ significantly between the open (ssDNA-bound) and closed (apo form) states, with the closed A3Bctd active site characterized by the distance between R211-Y315 as being less than 10 Å (Figure 2A). Additionally, we characterized the conformation of Y315 by its Chi1 dihedral angle, defining a closed conformation at 60° and an open conformation at 180° (Figure 2A).

**Figure 2.**
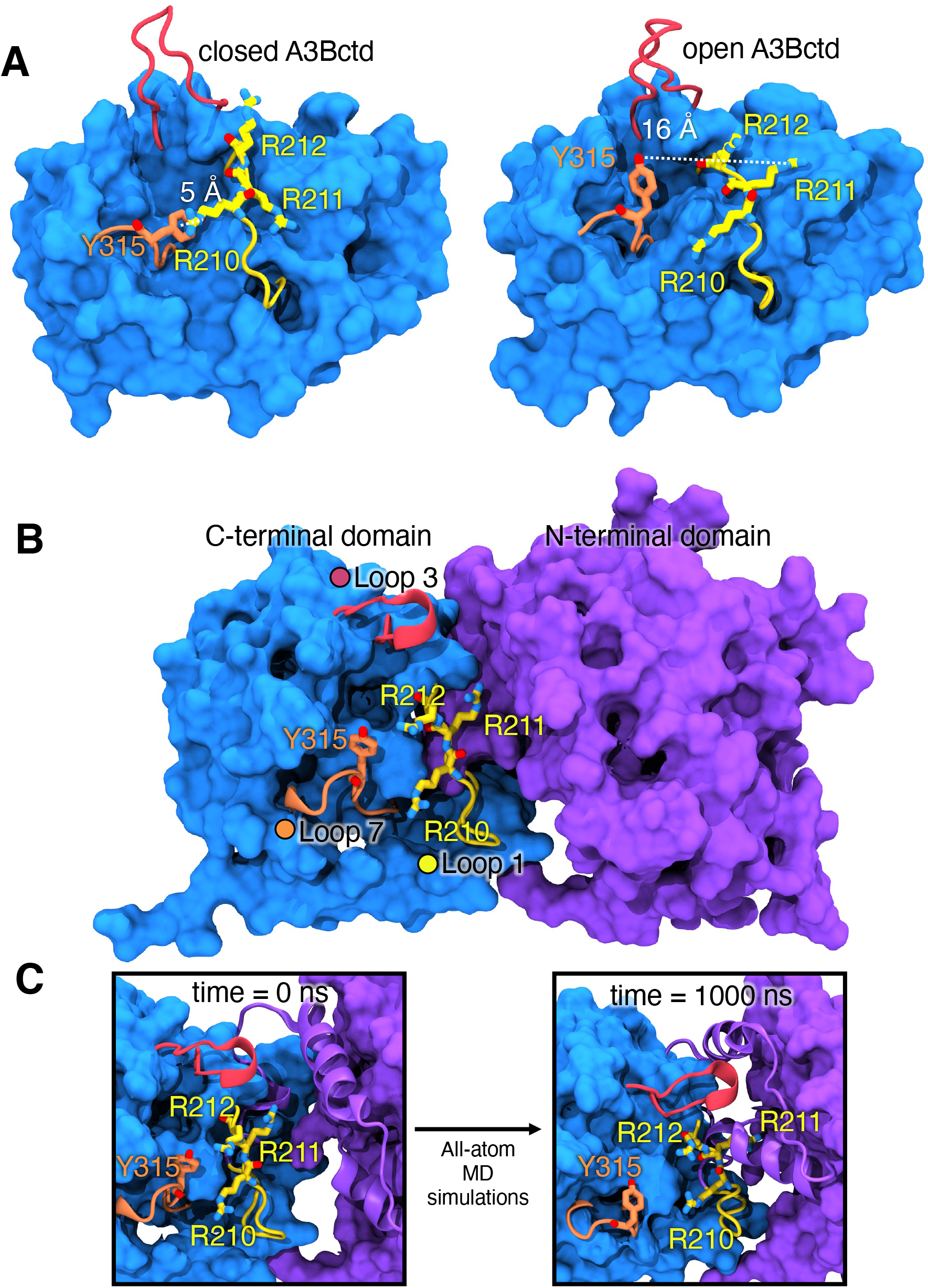
A3Bctd X-ray structures and fl-A3B model. (A) Representative structures of open and closed-state A3Bctd from available PDB X-ray structures (with ssDNA bound, not shown, from PDB ID 5TD5).^3^ The closed A3Bctd active site is defined with a 5 Å distance between R211-Y315. An open A3Bctd active site is defined with 16 Å distance between R211-Y315. (B) fl-A3B model. (C) Conformational dynamics of fl-A3B ssDNA active site, in loop 1, residues R210, R211, and R212 interacting with N-terminal domain.

The fl-A3B model structure used as starting point for MD simulations has a partially open active site, with the R211-Y315 interaction exhibiting a distance >10 Å and the Y315 Chi1 dihedral angle in trans conformation (180 degrees). This is because the template used for model construction was the ssDNA-bound A3Bctd discussed above. The fl-A3B was prepared for simulation (see Methods), with all missing loops restored, mutated residues reverted back to wild type, and explicit solvent and counter ions added. We subsequently performed 4 replicates of 1 μs conventional MD (4 μs aggregate sampling) for the fl-A3B model.

Overall the fl-A3B model exhibited stable dynamics over all 4 μs (SI Figure 3). In contrast with the A3Bctd, the fl-A3B active site opens more frequently (Figure 3), as exhibited by the shift to open-state for the R211-Y315 distance. Notably, in the A3Bctd alone, the R211-Y315 distance peaked near 10 Å, whereas in the fl-A3B, the R211-Y315 distance metric samples a significantly more open state, with distances ranging from 15 - 22 Å. The Y315 Chi1 dihedral angle in fl-A3B sampled both closed and open states, although it predominately sampled angles corresponding to the closed state.

**Figure 3.**
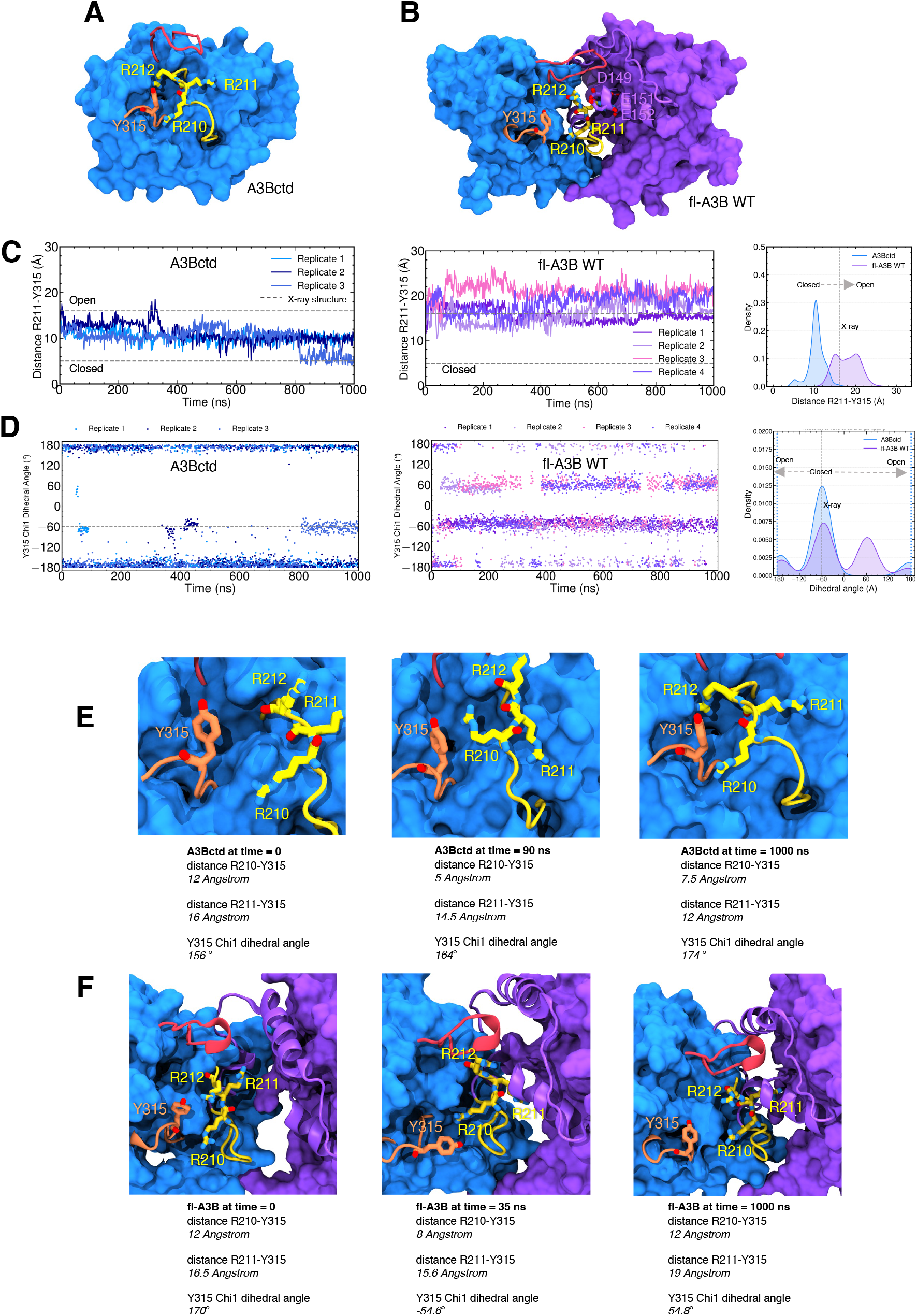
Structures and conformational dynamics of A3Bctd and fl-A3B. (A) A3Bctd; (B) fl-A3B; (C) fl-A3B and A3Bctd time-series distance of R211-Y315 with corresponding kernel density estimate (KDE) from several replicates of our simulations; (D) Time-series Chi1 dihedral angle; (E) A3Bctd representative snapshots highlighting the interactions of loop 1 residues R210, R211, and R212 with loop 7 residue Y315. Trajectories from our previous work were reanalyzed.^15,21^ (F) fl-A3B representative snapshots highlighting the interaction of R210, R211, and R212 with NTD residues.

To increase the likelihood of sampling the closed-to-open transition, we employed Gaussian accelerated molecular dynamics (GaMD) simulations in AMBER pmemd.cuda. GaMD, an enhanced sampling technique, uses a harmonic boost potential that smoothens energy barriers on the potential energy surface.^34^ During GaMD, the fl-A3B Y315 Chi1 dihedral angle values remained in the closed state (-60 degrees). The R211-Y315 distance is stable in its closed state at 9 Å (SI Figure 5). Thus, GaMD did not help in the opening transition of the fl-A3B active site.

To more rigorously characterize the active site opening mechanism, we subsequently carried out weighted ensemble (WE) simulations. WE is a transition pathway method that increases the chance of sampling biologically ‘rare’ events (e.g., in this case, sampling the closed to open active site transition).^35,36^ Application of this method requires selecting a so-called ‘progress coordinate’, i.e., a set of principle components that captures the critical dynamics of the system, to accurately describe the structural dynamics of interest in APOBEC3B. The optimal progress coordinate should reflect the slowest motions pertinent to the rare-event processes under investigation.^37^ Here, we used the R211-Y315 distance and the Y315 Chi1 dihedral angle as a 2-dimensional (2D) progress coordinate, where the closed state is defined by a R211-Y315 distance < 16 Å and a Y315 Chi1 dihedral of -50 to -70 degrees, and the target open state is defined by R211-Y315 distance > 16.0 Å and a Y315 Chi1 dihedral angle of -180 to 180 degrees.

Using the Python library WE Simulation Toolkit with Parallelization and Analysis (WESTPA) 2.0, we ran WE for both the A3Bctd and fl-A3B.^37^ We ran thousands of short MD simulations (0.05 ns) in parallel while iteratively replicating trajectories that transitioned from the closed to open state. Furthermore, a minimum adaptive binning (MAB) scheme was applied.^38^ Here, we maximized the successful trajectories by feeding back these trajectories to the initial state (i.e., steady state weighted ensemble).^39^ WE increased the sampling of the fl-A3B active site opening without altering the system’s free energy. The closed active site conformations derived from the fl-A3B GaMD simulations served as the initial structures for subsequent WE simulations. We simulated three independent replicates of fl-A3B closed-to-open state transitions to rigorously sample this active site opening.

Using the 2D progress coordinate, we observed over 7000 closed-to-open transitions leading to the active site opening of both A3Bctd and fl-A3B (Figure 4). As illustrated in Figure 4, including the NTD in the simulations increased the frequency of the fl-A3B active site opening. WE simulations demonstrated the efficient sampling of active site opening for A3Bctd and fl-A3B. Here, we observed that both models sampled the open states (i.e., -180.0 and 180.0 degrees). With Y315 Chi1 dihedral angle, the lowest energy minima accounted were at closed state, -60.0 degree and at open state -180 and 180 degrees. The wide distribution of R211-Y315 distance for the fl-A3B value was also observed. This is in contrast with the A3Bctd which is mostly concentrated at around 6.0 Å. Furthermore, in fl-A3B closed state Y315 Chi1 dihedral angle, we identified how the phenyl side chain blocked the ssDNA site (Figure 4C). The R211-Y315 distance distribution is much higher in fl-A3B than in A3Bctd. The fl-A3B R211-Y315 distance even reached >19.0 Å (Figure 4D). The low values of the free energy of closed-to-open state transition in WE were due to the lack of convergence. In both all-atom MD and WE method simulations, we observed a more frequent opening of the ssDNA active site in fl-A3B than in the A3Bctd alone.

**Figure 4.**
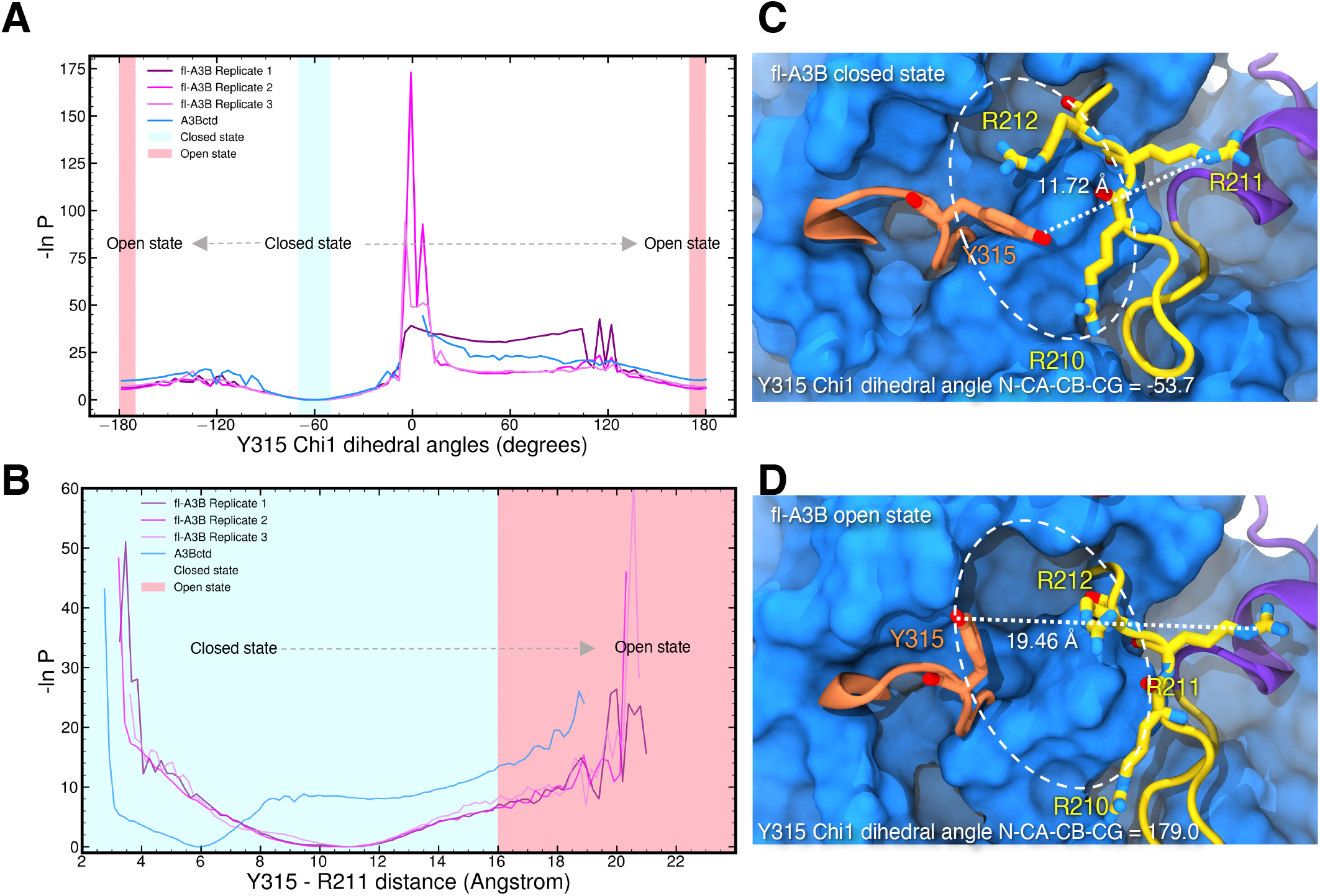
APOBEC3B ssDNA active site opening from steady-state weighted ensemble simulations. One-dimensional progress coordinate (A) Y315 Chi1 dihedral angle; (B) Y315-R211 distance; Probability distribution is plotted in an inverted natural log scale with free energy unit expressed as *1/k_BT_*. Transition of the closed state (blue region) to open state (red region) active site was marked with gray arrow. WE simulations of fl-A3B closed-to-open active site transitions were run in 3 replicates while A3Bctd with 1 replicate. (C) fl-A3B closed state; (D) fl-A3B open state. Active site was enclosed in an oval dashed line along with the corresponding Y315 Chi1 dihedral angle and R211-Y315 distance.

**Figure 5.**
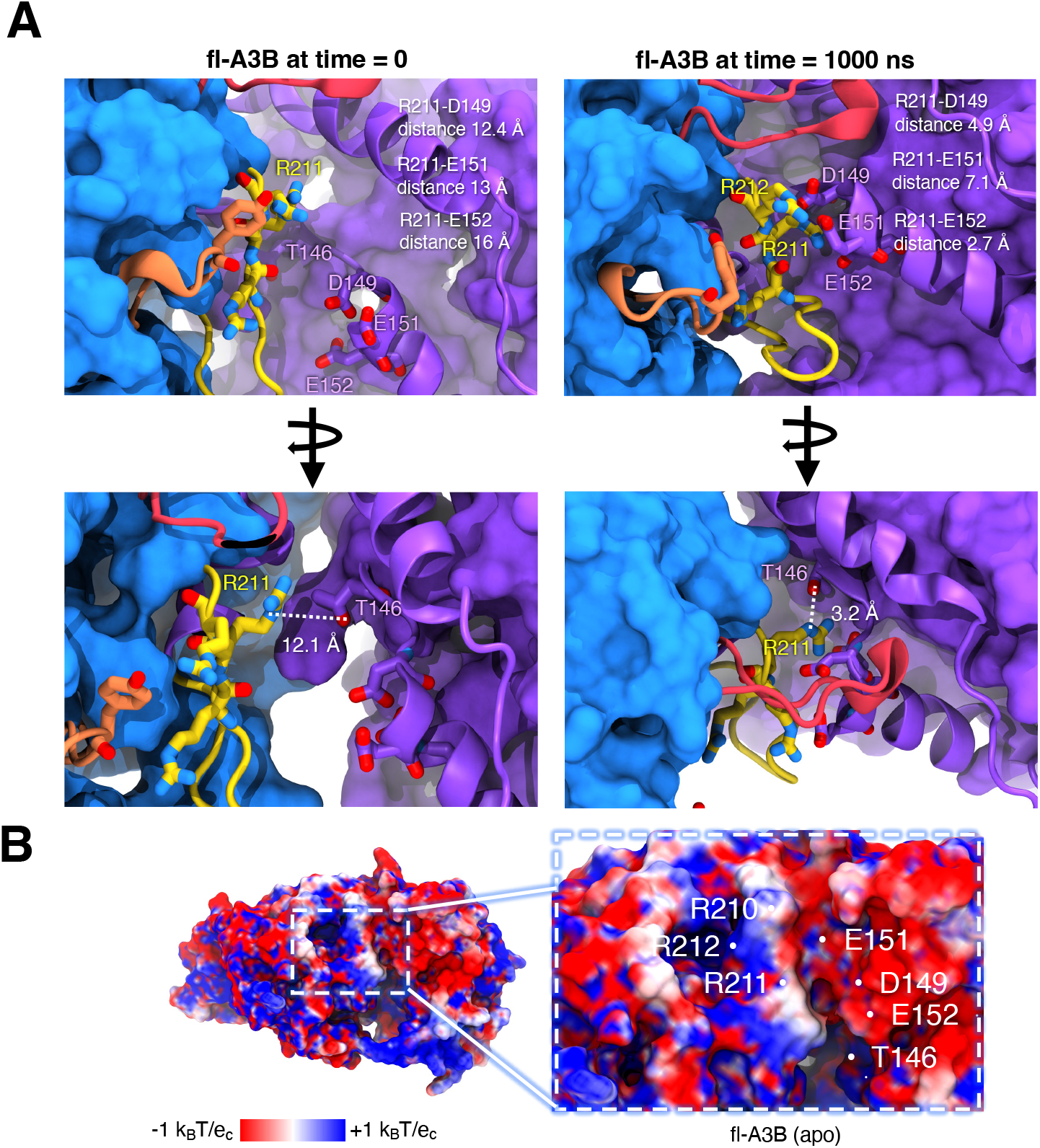
fl-A3B conformational dynamics showing the interactions of active site loop 1 residues with the NTD residues. (A) Interactions before and after molecular dynamics of the residues R210, R211, R212 with T146, D149, E151, and E152. (B) Adaptive Poisson-Boltzmann Solver (APBS) calculation on fl-A3B rendered in the top most clustered conformation from simulations.

### The N-terminal domain residues T146, D149, E151, and E152 form salt bridges with R210, R211, and R212

We hypothesized that the NTD of fl-A3B plays a crucial role in molecular recognition and stability, as evidenced by the binding and interactions of longer ssDNA (>10 nucleotides) with the NTD.^40^ In the apo form of fl-A3B, the residues T146, D149, E151, and E152 (in the NTD) form electrostatic salt bridges with residues R210, R211, and R212. We illustrated these interactions during 1 *μ*s simulations, showing a closer distance at the domain interface that impacts ssDNA binding (Figure 5A).

We used Adaptive Poisson-Boltzmann Solver (APBS) to visualize the electrostatic of the fl-A3B.^41^ Individual APBS calculations from multiple clustered structures from our MD simulations were averaged to consider the structural dynamics of the fl-A3B systems. In APBS calculations of the apo form and ssDNA bound fl-A3B, there is a highly negative patch at the NTD-CTD interface (Figure 5B). Interestingly, the positive charges in the side chains of R210, R211, and R212 can be attracted by the negative charges at D149, E151, and E152 allowing the favorable opening of the ssDNA active site due to a strong electrostatic interaction. This structural feature aligns with observed differences in enzymatic activity between fl-A3B and A3Bctd.

### Presence of the APOBEC3B N-terminal domain makes a more active enzyme

To assess relative rates of fl-A3B and A3Bctd deamination activity, we used purified recombinant proteins from human Expi293F cells and quantified activity using a recently developed real-time APOBEC3-mediated DNA deamination (RADD) assay.^42^ The RADD assay provides a direct read-out of APOBEC activity by including excess endonuclease Q (EndoQ) in each reaction, which functions downstream of C-to-U deamination to cleave newly uracilated substrate ssDNA oligonucleotide and thereby liberate fluorescence signal. Each enzyme was compared to a baseline oligo control reaction. A3Bctd showed nearly 50% lower deaminase activity compared to fl-A3B, as observed in real-time fluorescence readings (slope: 4.340 and 7.146, respectively) (Figure 6).

**Figure 6.**
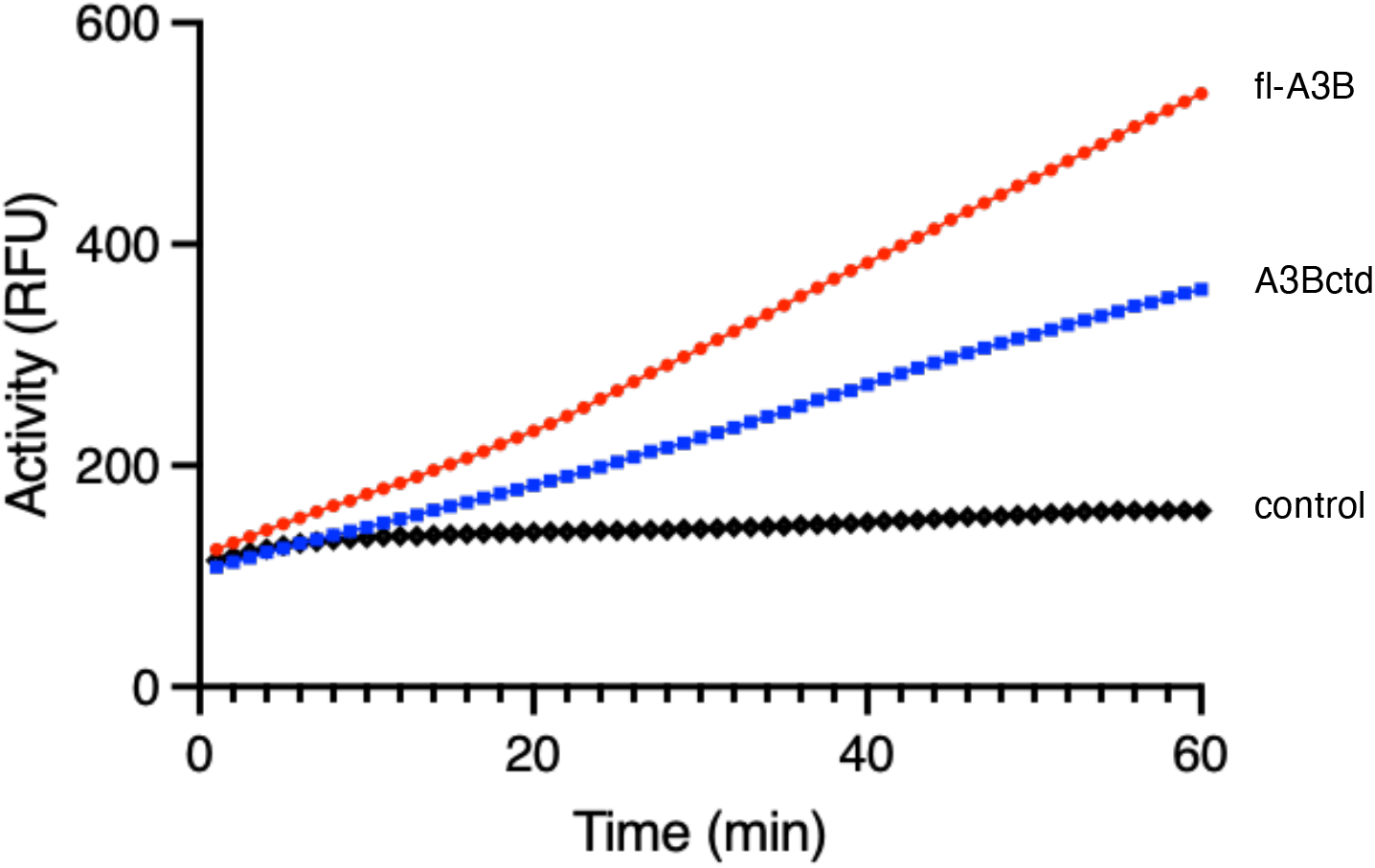
RADD assay readout for fl-A3B and A3Bctd. The fluorescent signal of 100 nM fl-A3B (red) and A3Bctd (blue) for a 1-hour reaction containing 0.8 μM reporter oligo is shown. The negative control (black) for the reaction contains 0.8 μM reporter oligo (mean fluorescent signal shown for n = 3 biological replicates).S

fl-A3B exhibits significantly higher activity and stability compared to A3Bctd as from previous experimental work. This aligns with the findings from Belica et al., who noted enhanced activity in both wild-type fl-A3B and A3Bctd,^42^ as well as with reports from Papini et al., which highlight the NTD’s role in allosteric regulation during the deamination reaction in longer ssDNA sequences.^40^ Kinetic analyses reveal that fl-A3B has comparable *K*_*m*_and higher *K*_*cat*_ than A3Bctd, underscoring its improved catalytic efficiency. Furthermore, in *Escherichia coli*, the expression and activity of fl-A3B exceed those of A3Bctd by four-to eight-fold.^43^ Further corroborating these functional enhancements, Butt et al. reported that the NTD not only accelerates substrate search but also contributes to the overall enzyme stability.^43^ This is despite fl-A3B and A3Bctd having similar preferences for structured ssDNA targets, such as hairpin loops over linear DNA forms.

The NTD of fl-A3B is crucial for its biological function, influencing both cellular localization and ssDNA interactions.^44,45^ Key residues within the NTD, such as Y18 and D19, are essential for nuclear localization, and W127 is important for ssDNA binding and regulation, as mutations in these residues lead to reduced catalytic efficiency.^44^ The NTD also enhances the enzyme’s functional stability and interaction with ssDNA, supported by its role in forming high molecular weight complexes and its involvement in RNA-dependent attenuation processes.^14^ This evidence synergistically provides a more comprehensive view of how the NTD can have multiple roles in ssDNA deamination, activation, allosteric inhibitor recognition, and nucleic acid substrate processivity.

Overall, we surmised how the NTD can increase the activity of fl-A3B and its unique interaction with the CTD, which has implications for several cancers in terms of inhibitor discovery and design. This multifunctional domain thus plays a pivotal role in the dynamical and functional properties of fl-A3B, highlighting its significance beyond nucleating RNA interactions.

## Summary and Conclusion

A long-term goal for fl-A3B is to fully characterize the thermodynamics, kinetics, and structural and dynamic properties to better inform drug discovery efforts against various cancer types. This study employed multi-microsecond, all-atom, explicitly solvated MD simulations, including conventional, GaMD, and WE simulations, to explore the conformational dynamics of fl-A3B with a particular focus on the active site opening. Our results demonstrate that the N-terminal domain (NTD) significantly enhances the likelihood of active site opening in fl-A3B through salt bridge interactions between active site residues R210, R211, R212, and NTD residues T146, D149, E151, and E152. Along with the available experimental results, we showed that the interaction of the CTD and NTD in the fl-A3B facilitate the active site opening, thus making the ssDNA deamination more efficient. The insights gained from this mechanistic exploration inform our general understanding of DNA editing systems and may improve attempts to develop more selective APOBEC inhibitors.

## Systems and Methods

In summary, we built a complete full-length A3B model based on the known A3Bntd and A3Bctd X-ray structures. This model was compared with cryo-EM micrographs of the fl-A3B. After validating our model, we ran all-atom MD simulations (i.e., conventional, GaMD, and WE simulations) to sample structural changes in the two domains and the ssDNA active site. Structural determinants of active site opening were measured, analyzed, and complemented by A3B deamination assays (i.e. RADD).

### Full-length APOBEC3B model generation

To provide a 3D model of the wild-type A3Bctd, we reverted the crystallized ssDNA-bound A3Bctd construct (PDB ID 5TD5)^10^ back to wild-type (WT) using *in silico* modeling tools as described in our previous work.^15,21^ SI Figure 1 outlines our fl-A3B model construction. Using the same protocol, the structure of an A3Bntd construct (PDB ID 5TKM)^14^ was also reverted back to wild-type. The ClusPro web server was then used to generate 30 protein-protein docked poses of A3Bctd and A3Bntd.^46^ Filtering out the poses that had unreasonable linker distance between the N-terminal residue of A3Bctd and the C-terminal residue of A3Bntd left us with three docked poses. We then extended the missing 6 residues at the N-terminus of A3Bntd and the missing 3 residues at the C-terminus of A3Bctd using Chemical Computing Group’s MOE molecular modeling tools for all 3 docked poses selected.^47^ Finally, we used Schrodinger’s Prime module to generate 3 homology models of fl-A3B with the sequence from UniProt (ID Q9UH17) using the 3 docked poses as templates.^24^ The closest model to the cryo-EM micrograph data was used in subsequent MD simulations.

### Conventional all-atom molecular dynamics (MD) simulations

All-atom MD simulations were run on the Delta supercomputing system at the National Center for Supercomputing Applications (NCSA) using AMBER’s GPU MD engine pmemd.cuda.^29^ Our MD simulation workflow was consistent with the previous work reported.^15,21^ Four replicates for each of the systems were simulated. AMBER ff14SB and TIP3P force fields were used for protein and water, respectively.^30,31^ VMD PROPKA was used for assigning protein protonation states at pH 7.^48^ The Zn^2+^ ion was modeled with the Cationic Dummy Atom Model.^49^ Each system underwent energy minimization, followed by gradual heating, and equilibration with decreasing restraints. In the first 500 steps of minimization, only hydrogen atoms were minimized leaving all other atoms fixed. In the second 500 steps, all hydrogen atoms, water molecules, and ions (except Zn^2+^ ion) were minimized. In the third 500 steps, all atoms except the protein backbone were minimized. The entire systems were then minimized without any restraints for 40,000 steps. After the minimizations, each system was slowly heated up from 0 K to 310 K using a positional restraint of 2.0 kcal/mol*/*Å^2^ on all non-hydrogen atoms of protein except the zinc-coordinating residues. Each system was then relaxed further in three consecutive 250 ps restrained MD simulations at 310 K and 1 atm using restraints of 3.0, 2.0, and 1.0 kcal/mol*/*Å^2^ on all non-hydrogen atoms of protein except the zinc-coordinating residues. In total, we gathered 4 replicates of one microsecond, totaling 4 μs MD simulation data. We also used the apo form A3Bctd MD simulations datasets we previously reported for comparison.^15,21^

### Gaussian accelerated molecular dynamics (GaMD) simulations

After this multi-step system relaxation protocol, GaMD simulations were performed without any restraints.^34^ A 2 fs time step was used. Three replicates of GaMD were run based on the three models of fl-A3B generated. Temperature was maintained at 310 K using Langevin dynamics with a collision frequency of 5 ps and pressure was maintained at 1 atm using isotropic position scaling with a pressure relaxation time of 2 ps. Long-range electrostatics was treated by the Particle Mesh Ewald method, and a non-bonded cutoff of 10 Å was used.^29^ The SHAKE algorithm was used to constrain the bonds involving hydrogen atoms.^50^ Given an average system size for protein, the maximum, minimum, average, and standard deviation values of the system potential have been obtained from an initial 1.2 ns NPT simulation with no boost potential.

Finally, 300 ns of GaMD simulations have been carried out. Each system was simulated in 3 replicates, differing in the initialization seed for the velocities, and each replicate underwent 300 ns of unrestrained MD simulation in an NPT ensemble.

### Weighted ensemble (WE) method simulations

WE Simulation Toolkit with Parallelization and Analysis (WESTPA) 2.0, a software package in Python and a tool for constructing and running stochastic simulations with the weighted ensemble method approach, was utilized.^37,51,52^ To initialize the WE simulations, we employed 100 closed fl-A3B active site structures, each assigned equal statistical weights of 0.01. Progress coordinates (fl-A3B Y315 Chi1 dihedral angle and the R211-Y315 distance), which define the closed and open active site states, were calculated using CPPTRAJ.^53^ A Y315 Chi1 dihedral angle is considered closed or open if the values -60 and 180 degrees are reached, respectively. For tyrosine residue, the Chi1 dihedral angle is defined as the angle between atoms N-CA-CB-CG where the CG *cis* to N is defined as a 0-degree angle. The target open state, defined by a Y315 Chi1 dihedral angle of -180 to 180 degrees and an R211-Y315 distance of at least 16.0 Å, was continually sampled. Trajectories that reached the open state in a non-equilibrium steady state were recycled, initiating new trajectories from the initial state with carried-over trajectory weight to enhance sampling of close-to-open active site transitions.^39^Non-equilibrium steady-state WE simulations were conducted to sample transitions from the closed to open active site states efficiently.

A short simulation time of 50 ps, denoted as *τ*, was implemented for resampling using the Minimum Adaptive Binning (MAB) scheme.^38^ In this scheme, the positions of bins are strategically placed along the defined progress coordinates (i.e., Y315 dihedral angle and R211-Y315 distance) and adjusted according to trailing and leading trajectories using the westpa.core.binning module specified in the west.cfg file.

Our simulations were executed at the National Center for Supercomputing Applications (NCSA) Delta supercomputer utilizing NVIDIA A100 GPUs. Dynamics were propagated using AMBER pmemd.cuda with the temperature maintained at 300 K using a Langevin thermostat and pressure controlled by a Berendsen barostat within an orthogonal box.^29^ The fl-A3B closed-to-open state non-equilibrium steady state weighted ensemble simulations were conducted in triplicates, accumulating a total simulation time of 45.58 μs data.

### MD simulations data analysis

Distances between the center of mass of residue were measured using pytraj.^53^ The average potentials from these clusters were plotted back to the structure of fl-A3B. The root mean square deviation (RMSD) of the protein backbone atoms (C, CA, N, and O) was calculated with pytraj, using the first frame of the production run as a reference. To account for global translational and rotational movements in the RMSD calculations, all frames were fitted to the reference frame. A combined histogram of these RMSD values was also generated. Visualization of all structures and trajectory movies was done using VMD.^54^ Clustering based on the RMSD of both R211 and Y315 Cα was done using the Python package scikit-learn’s kmeans method.^55^ The clustered structures were also subjected to Adaptive Poisson Boltzmann Solver (APBS) electrostatic calculations using the APBS package (version 3.0).^41^

### fl-A3B protein expression and purification for cryo-EM data collection

The A3B protein used in cryoEM studies contained human A3B residues 7 to 378 fused to maltose-binding protein (MBP) on the N-terminus and 6xHis-tag on the C-terminus. This A3B construct had solubility-enhancing mutations (Y13D, F61S, Y65H, Y83D, W127S, T146K, Y162D, F200S, W228S, L230K, F308K) and an internal truncation of loop 3.^16^ The protein was expressed in *E. coli* strain BL21(DE3) and purified by nickel-affinity and size-exclusion chromatography as a monomer. Purified protein in 20 mM Tris-HCl (pH 7.4), 0.5 M NaCl, and 5 mM 2-mercaptoethanol was flash-frozen and stored at –80°C. The protein concentration was determined based on UV absorption at 280 nm.

### CryoEM data collection

Purified fl-A3B, prepared at 1 mg/mL, was deposited (3 µL) onto UltrAuFoil R1.2/1.3 300-mesh grids, which were cleaned for 10 seconds using a Solarus plasma cleaner (Gatan, Inc.) at 15 watts in a 75% nitrogen/25% oxygen atmosphere. The sample was manually blotted for 6-7 seconds with Whatman No. 1 filter paper and immediately plunge-frozen in liquid ethane, cooled by liquid nitrogen, under cold room conditions (^3^ 95% humidity at 4°C). Squares were selected for the thinnest ice.

Micrographs of the A3B complex were acquired using a Talos Arctica electron microscope (Thermo Fisher Scientific) operating at 200 kV, equipped with a K2 Summit direct electron detector (Gatan, Inc.). A total of 1,420 micrographs were recorded. Movies of A3B were captured in counting mode at a pixel size of 1.16 Å/pixel and a magnification of 36,000x, with a defocus range between −1.7 µm and −2.1 µm. The acquisition was performed using an exposure rate of 6.98 e^-^/pixel/s over 12.5 seconds (250 ms per frame, 50 frames), resulting in a total dose of approximately 65 e^−^/Å^2^ (1.32 e^−^/Å^2^ per frame). Patch-based motion correction and contrast transfer function (CTF) estimation were carried out in cryoSPARC Live.^56^ Particle picking was performed using the Blob Picker, with particle diameters ranging from 50 Å to 200 Å. A total of 3,417,871 particles were extracted with a box size of 256 pixels. One round of 2D classification was conducted using 50 classes, an initial uncertainty factor of 2, and an inner mask diameter of 240 Å. An ab initio reconstruction was generated from 100,000 particles, with an initial resolution of 35 Å and a maximum resolution of 12 Å. The final reconstruction achieved a resolution of 12 Å.

### fl-A3B and A3Bctd protein purification

Protein expression and purification as previously described with modifications.^57^ Briefly, using the standard Expi293 transfection protocol (ThermoFisher), 25 mL cultures of Expi293F cells (ThermoFisher) were transfected with pcDNA3.1-hA3Bi-mycHis or pcDNA3.1-A3Bctd (193-382)-mycHis. Transfection enhancers were added 18-20 hours post-transfection. Cells were harvested 3 days post-transfection by centrifugation at 1,000 rpm for 5 minutes; then the cell pellets were frozen at -80*°*C. Cells were thawed on ice, then resuspended in 15 mL lysis buffer (25 mM Tris-Cl pH 8.0, 500 mM NaCl, 5 mM MgCl_2_, 20 mM imidazole, 5% glycerol, and 0.1% IGEPAL CA-630), followed by sonication on ice for 2 minutes, at power 5 and 40% duty. RNase was added to 100 *μ*g/mL, as well as 100 U of Salt Active Nuclease (MilliporeSigma), and incubated at 37*°*C for 2 hours for nucleases to act. Following centrifugation (16,000 x g, 30 minutes), clarified lysates were collected, and sodium chloride was added to a final concentration of 1 M. 100 *m*L Ni-NTA Superflow resin (Qiagen) was added to the lysates and incubated for 1 hour at 4*°*C with agitation. The resin was washed with 1 CV wash buffer (25 mM HEPES pH 7.4, 0.1% Triton X-100, 10% glycerol, 300 mM NaCl and 40 mM imidazole) and then incubated 3 times with 200 *μ*L of an elution buffer (25 mM HEPES pH 7.4, 0.1% Triton X-100, 20% glycerol, 150 mM NaCl, 400 mM imidazole, and 1 mM TCEP) for 10 minutes to elute proteins. Elutions were then pooled and concentrated using an Amicon (MilliporeSigma) device with a 10 kDa cut-off before storage. Proteins were snap-frozen in liquid nitrogen and stored in 5 *μ*L aliquots at -80*°*C. Once thawed, proteins were kept on ice for no longer than 24 hours and never refrozen.

### *Pyrococcus furiosis* Endonuclease Q purification

BL21(DE3)-Star cells were transformed with pET24a-PfuEndoQ-6xHis.^58^ PfuEndoQ was purification was adapted from previous works.^58,59^ Briefly, single colonies were picked into an overnight starter culture in Terrific Broth, which was used to inoculate 1 L of Terrific Broth. Cells were grown at 37°C until the OD600 reached ∼1. Expression of Pfu EndoQ was induced by the addition of 0.5 mM IPTG followed by shaking overnight at 18°C. Cells were harvested by centrifugation and lysed by addition of 100 mL of 50 mM Tris-Cl pH 8.0, 10% glycerol, 600 mM NaCl, 1 mM MgCl_2_, two Roche cOmplete EDTA-free Protease Inhibitor Cocktail tablets (MilliporeSigma), and hen egg white lysozyme to 1 mg/mL followed by rocking for 30 minutes. The lysate was then sonicated with a Branson Sonifier 450 for 4 two-minute cycles at 60% duty cycle, power 6, with at least 1 minute between runs. Imidazole was added to 20 mM and TCEP to 0.5 mM and the lysate was heated to 70°C for 30 minutes to denature most of the *E. coli* proteins. The lysate was then centrifuged for 30 minutes at 16,000 x g to pellet insoluble material. The resulting clarified lysate was purified over a two mL Ni-NTA column (Qiagen 30230), washed twice with 5 column volumes (CV) of 50 mM Tris-Cl pH 8.0, 300 mM NaCl, 20 mM imidazole, and eluted in 12 mL of 50 mM Tris-Cl pH 8.0, 150 mM NaCl, 400 mM imidazole, 0.5 mM TCEP. Following analysis by SDS-PAGE, the protein was dialyzed into storage buffer (25 mM Tris-Cl pH 8, 150 mM NaCl, 50% glycerol) and stored at -20°C with 0.5 mM TCEP.

### Real-time APOBEC3-mediated DNA deamination (RADD) assay

Each 20 *μ*L reaction was carried out on a flat-bottom black 384-well plate (Corning Ref. 3821). Prepared enzyme mixture, consisting of 200 nM A3B (A3Bfl, A3Bctd or control) in 1x NEBuffer 2 (New England Biolabs) made in nuclease-free water and oligo mixture, consisting of 4 *μ*M EndoQ and 1.6 *μ*M TC oligo (5’-6-FAM-TAGGT**<u>C</u>**ATTATTGTG-IAB-3’) in 1x NEBuffer 2 made in nuclease-free water. Added 10 *μ*L of enzyme mixture to each well, followed by 10 *μ*L of oligo mixture (final concentrations for A3B, EndoQ, and substrate oligonucleotide are 100 nM, 2 *μ*M, and 0.8 *μ*M, respectively). All fluorescence readings were recorded instantly on a TECAN Spark 10M instrument in concert with the SparkControl V3.0 software. Data was collected every minute for 1 hour at 37*°*C using the scanning fluorescence kinetic mode (excitation: 485 nm; emission: 535 nm) with the gain set to 50%. Slope determination was performed by simple linear regression using Prism 10.

## Abbreviations

A3A: APOBEC3A
A3B: APOBEC3B
A3Bctd: APOBEC3B C-terminal domain
A3Bntd: APOBEC3B N-terminal domain
A3G: APOBEC3G
APOBEC3: Apolipoprotein B mRNA editing enzyme, catalytic polypeptide-like 3
CTD: C-terminal domain
fl-A3B: Full-length APOBEC3B
GaMD: Gaussian accelerated molecular dynamics
MD: molecular dynamics
NTD: N-terminal domain
RADD: Real-time APOBEC-mediated DNA deamination
RFU: Relative fluorescence unit
UDG: uracil-DNA glycosylase
WE: Weighted ensemble method

## Acknowledgments

Authors would like to thank Amaro Lab members, especially Mohamed Shehata and Fiona Kearns for their input in the improvement of the computational workflow and data analyses. M.K.E.B. and S.-H.A. would like to thank Lillian Chong and their group for their help in running weighted ensemble simulations. The authors would like to thank Nuri Alpay Temiz, Geoffrey Jameson, and Daniel Harki for helpful feedback. S.-H.A. would like to acknowledge the University of California, Davis for the start-up funds. C.K.J.M. was supported by an NIH, NCI traineeship (T32-CA009523). C.C.T. was supported by Schmidt Sciences, LLC. This work was supported by NCI P01-CA234228 to R.S.H., H.A., and R.E.A., NIGMS R35-GM118047 to H.A., and a Recruitment of Established Investigators Award from the Cancer Prevention and Research Institute of Texas (CPRIT RR220053 to R.S.H.). Supercomputing resources are provided by XSEDE (NSF TG-CHE060073) and ACCESS (NSF TG-CHE060063) allocations granted to R.E.A. R.S.H. is the Ewing Halsell President’s Council Distinguished Chair at University of Texas San Antonio and an Investigator of the Howard Hughes Medical Institute.

## Conflict of Interest

The authors declare no conflict of interest.

## Authors Contribution

R.E.A., H.A., and R.S.H. designed and oversaw the research project and secured funding. M.K.E.B. and Ö.D. built the full-length APOBEC3B model and performed MD simulations. M.K.E.B., Ö.D., S.-H.A., and C.K.M. performed weighted ensemble simulations. M.K.E.B., Ö.D., S.-H.A., C.K.M., C.C.T., L.C., and R.E.A. performed simulation analyses and data visualizations. M.A.C., Y.C., B.d.l.P.A., R.S.H., and H.A. run protein biochemistry and purification experiments. K.L.M. and M.A.H.,Jr. processed cryoEM dataset. M.K.E.B., Ö.D., and R.E.A. wrote the initial draft of the paper with contributions from all authors. All authors reviewed and approved the manuscript.

## Software and Data Availability

Trajectories, structures, simulation scripts, analysis scripts, and data files are uploaded in our group’s website and can be accessed thru this link https://amarolab.ucsd.edu/data.php. All software used is community developed and freely available.

## Notes

### Competing Interest Statement

The authors have declared no competing interest.

