## Supporting Information for "Regulatory interactions between APOBEC3B N- and C-terminal domains"

### SUPPLEMENTARY INFORMATION

Contents

Figures

**SI Figure 1**. CryoEM data processing of the full-length APOBEC3B and maltose binding protein complex.
**SI Figure 2**. Workflow in modeling and refining the human full-length APOBEC3B (fl-A3B).
**SI Figure 3**. Protein backbone root-mean-square deviation.
**SI Figure 4**. R210-Y315, R211-Y315, and R212-Y315 distances.
**SI Figure 5**. Representative conformation of the Y315 Chi1 dihedral angle in a partially open and open states active site.
**SI Figure 6.** GaMD simulations of fl-A3B.

Movies

**SI Movie 1**. A3Bctd dynamics from classical MD simulations.
**SI Movie 2**. fl-A3B dynamics from classical MD simulations.

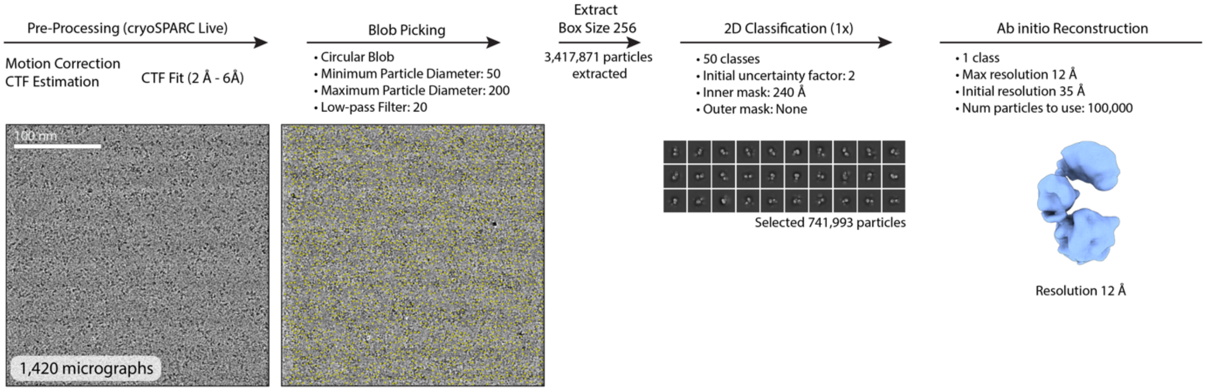


**SI Figure 1**. CryoEM data processing of the full-length APOBEC3B and maltose binding protein complex. See main text Methods section for detailed explanation of cryoEM data processing.

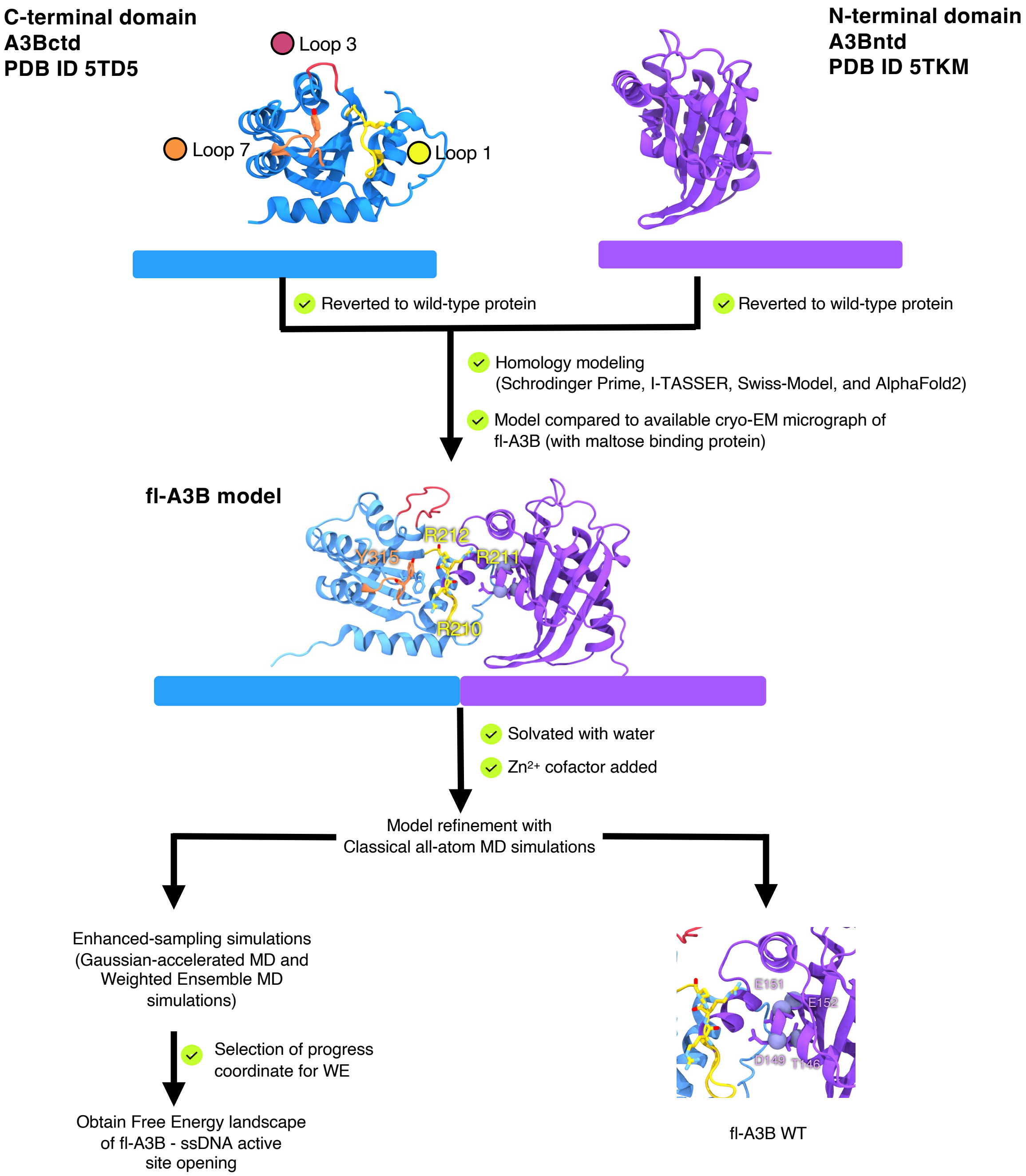


**SI Figure 2.** Workflow in modeling and refining the human full-length APOBEC3B (fl-A3B). Reference structures for homology modeling of fl-A3B PDB ID 5TD5 (ssDNA substrate removed)[3] and 5TKM have several mutations. We reverted them to the wild-type A3Bctd and A3Bntd. Models were refined with conventional all-atom MD simulations and were used for further modeling and simulations work (i.e., Gaussian accelerated and WE method MD simulations and mutations study on fl-A3B N-terminal domain residues).

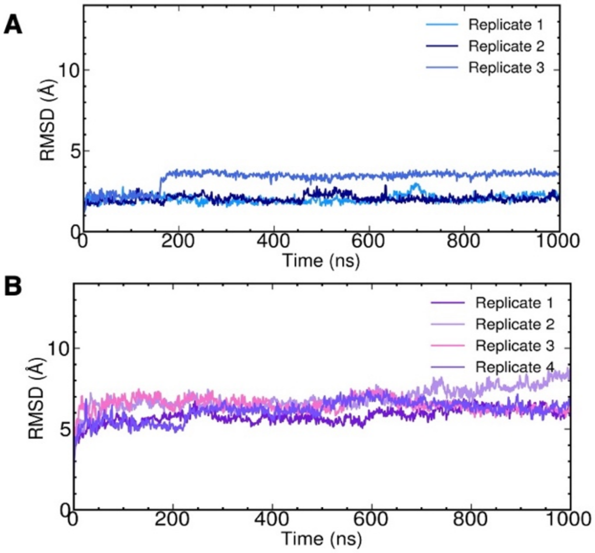


**SI Figure 3**. Protein backbone root-mean-square deviation (RMSD). (A) A3Bctd and (B) fl-A3B Wild-type.

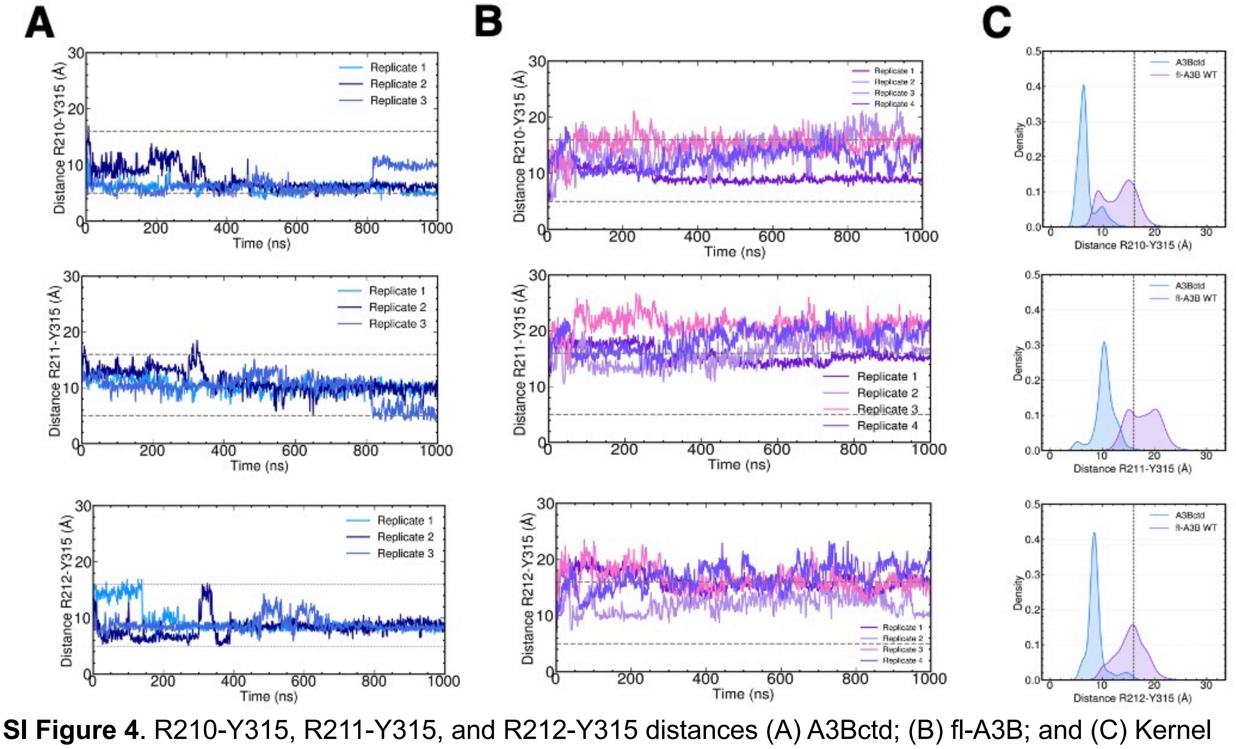


**SI Figure 4**. R210-Y315, R211-Y315, and R212-Y315 distances (A) A3Bctd; (B) fl-A3B; and (C) Kernel Density Estimates of the distances.

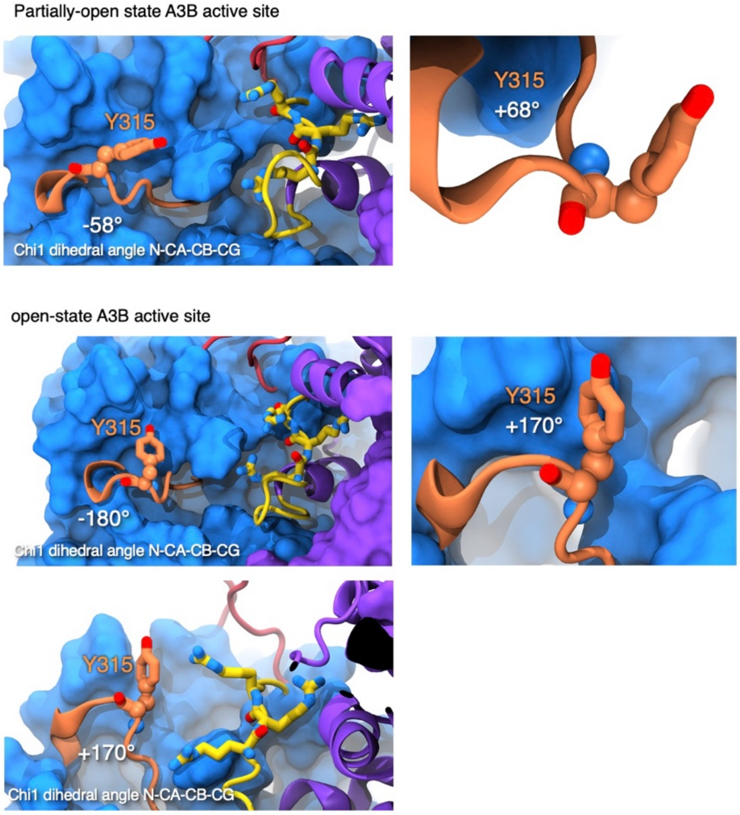


**SI Figure 5**. Representative conformation of the Y315 Chi1 dihedral angle in a partially open and open states active site.

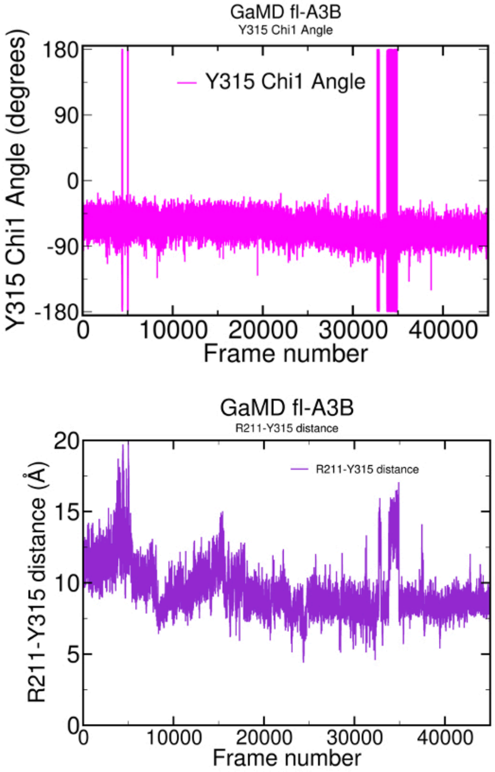


**SI Figure 6.** GaMD simulations of fl-A3B. Y315 Chi1 dihedral angle values (top) and R211-Y315 distance values (bottom).
